# Impact of fruit-tree shade intensity on the growth,yield, and quality of intercropped wheat

**DOI:** 10.1101/396010

**Authors:** Xu Qiao, Lihan Sai, Xingwu Chen, Lihua Xue, Junjie Lei

**Affiliations:** Institute of Grain Groups, Xinjiang Academy of Agricultural Sciences, Urumqi, Xinjiang 830091, China; Institute of Medicinal Plant Development, Chinese Academy of Medical Sciences, Beijing 100193, China

**Keywords:** Agroforestry system, Intercropping, Wheat, Photosynthetically active radiation, Quality

## Abstract

Agroforestry is a common traditional practice in China-especially in the southern Xinjiang of Northwest China. However, the productivity of many agroforestry systems has been lower than expected in recent years, highlighting the need for an actionably deep mechanistic understanding of the competition between crops and trees. Here, we chose 3 different fruit tree/wheat intercropping agroforestry systems to investigate influence of different fruit tree shade intensity on the growth, yield and quality of intercropping wheat: jujube/wheat, apricot /wheat, and walnut /wheat. We found that compared to the monoculture wheat system, the mean daily shade intensity of the jujube-, apricot-, and walnut-based intercropping systems were, respectively, 23.2%, 57.5%, and 80.7% shade. The photosynthetic rate of wheat in the jujube-, apricot-, and walnut-based intercropping systems decreased by, respectively, 11.3%, 31.9%, and 36.2% compared to monoculture wheat, and the mean number of fertile florets per spike decreased by 26.4%, 37.4%, and 49.5%. Moreover, the apricot- and walnut-based intercropping systems deleteriously affected grain yield (constituent components spike number, grains per spike, and thousand grain weight) and decreased the total N, P, and K content of intercropping wheat. Tree shading intensity strongly enhanced the protein content, wet gluten content, falling number, dough development time, and dough stability time of wheat, but significantly decreased the softening degree. Strong negative linear correlations were observed between tree shade intensity and the number of fertile florets, grain yield (including spike number, grains per spike, and thousand grain weight), nutrient content (N, P and K), and softening degree of wheat. In contrast, we found that daily shade intensity was positively linearly correlated with protein content, wet gluten content, falling number, dough development time, and dough stability time. We conclude that jujube-based intercropping systems can be practical in the region, as they do not decrease the yield and quality of intercropping wheat. Further research should focus on the above-ground/below-ground interspecific interactions and the mechanisms behind the observations that we observed amongst the intercropping systems.

## Introduction

Agroforestry is a land-use system in which woody perennials are grown in association with agricultural crops or pastures, in which there are both ecological and economic interactions between trees and the other components [1]. Agroforestry systems are increasingly viewed as having significant potential to provide a range of environmental services, including reductions in nutrient leaching, improvements in soil erosion and water loss [2,3], enhancement of soil nutrient status and nutrient cycling [4], sequestration of carbon [5], increases in soil organic carbon, increases in soil microbial community diversity and abundance [6], and increases in the effects of the activity of beneficial soil organisms [7]. Additionally, agroforestry systems can provide windbreaks, thereby reducing wind speed [8]. Tree-based intercropping systems also promote larger earthworm populations compared to monoculture crops [9]. *Zizyphus jujuba*–*Triticum aestivum* agroforestry systems are frequently used to improve land-use efficiency and increase economic returns in southern Xinjiang Province [10].

Friday and Fownes [11] reported that competition for light is the main cause of reductions in maize yields in hedgerow/maize intercropping systems in the USA. Kittur et al. [12] reported that low understory photosynthetically active radiation (PAR) was the dominant factor contributing to reductions in the growth of turmeric in denser bamboo stands compared to widely spaced bamboo in India. Similar results were reported in *Paulownia* systems on the North China Plain and Loess Plateau [13-15]. Jose et al. [16] observed that maize yields were reduced by 35% and 33% when intercropped with black walnut and red oak, respectively, compared to monoculture treatments. Smethurst et al. [2] also found that competition for light was the main factor causing lower crop yields compared to a monoculture configuration in a temperate agroforestry system, and that C4 crops (e.g., maize) were more vulnerable to shading compared to C3 crops (e.g., wheat). Wang et al. [17] reported that the yields of both jujube and wheat were lower in 3-, 5-, and 7-year-old jujube tree–wheat intercropping systems, and that the wheat yield decreased as the distance from the jujube trees decreased. Thus, it is important to investigate the mechanisms of aboveground competitive interactions in agroforestry systems.

By 2012, the total area of fruit trees in southern Xinjiang Uygur Autonomous Region, Northwest China, had reached more than 1 million hectares [10]. Fruit tree-based intercropping systems are widely favored by the local population, and more than 80% of fruit trees have been planted as intercropping systems. However, as the fruit trees have grown, the productivity of the intercropping crops in many of the agroforestry systems has been lower than expected in recent years [10,15,17]. The widespread planting of fruit trees has consequences for food security, and has challenged the ability of the region to feed the local population. Therefore, it is important to highlight the need for a systematic understanding of belowground and aboveground interactions under different agroforestry systems to guide practices that can achieve high yields and efficiency.

Although many of the competitive pathways in alley cropping systems have been identified, not all have been adequately quantified. In this study, we compared three different varieties of fruit-tree (jujube, apricot, and walnut) intercropping with wheat to examine the aboveground interactions and likely response mechanisms. Fruit trees and wheat were selected for study because of their importance as the main economic and food crops in southern Xinjiang Uygur Autonomous Region, Northwest China. The objectives of the study were to determine (1) whether the fruit trees had a significant effect on the growth and yield of the companion crop (wheat) via shading; (2) whether the yield of the intercropped plants could be increased in this agroforestry system, and what possible solutions are available to minimize aboveground interspecies competition; (3) whether this planting mode is suitable, and which fruit tree-based intercropping system offers the best option in the region; and (4) the effects of this agroforestry system on the quality of the intercropped wheat.

## Materials and methods

### Site description

Field experiments were conducted in 2011 and 2012 at 4 Village of Zepu County (38°05′N, 77°10′E), Kashi Prefecture, Xinjiang Uygur Autonomous Region, China. Altitude is 1,318 m above sea level. Annual mean temperature is 11.6 °C (1961-2008). Cumulative temperatures above 0 °C is 4,183 °C. The mean frost-free period is 212 days. Annual precipitation is 54.8 mm, potential evaporation is 2,079 mm. This region has a typical arid climate, and the soil type is arenosol. Some chemical properties of the soil are presented in S1 Table.

### Experimental design

The experimental design for this study was a field experiment, comprising monoculture wheat (*Triticum aestivum* L. Xindong-20), wheat intercropped with 9-year-old jujube trees (*Zizyphus jujuba* Mill. Junzao), wheat with 10-year-old apricot trees (*Prunus armeniaca* L. Saimaiti), and wheat with 10-year-old walnut trees (*Juglans regia* L. Wen-185). The row distance was 0.13 m in wheat. The fruit trees were planted in north-south orientation. Basic information for the different types of fruit trees is showed in S2 Table. The jujube-, apricot-, and walnut-based intercropping wheat strips were, respectively, 3.30, 5.10, and 6.00 m wide, and the distance between the jujube, apricot, and walnut tree to the nearest wheat row was 0.85, 0.95, and 1.00 m; the jujube, apricot, and walnut tree occupied 34.0%, 27.1%, and 25.0% of the gross field site areas (Fig 1). The total area for each of the four systems (monoculture wheat, jujube-wheat, etc) was 0.4 hm^2^. The sown density of monoculture wheat and the 3 intercropping systems were each 4.25×10^6^plants per hm^2^.

**Figure 1.**
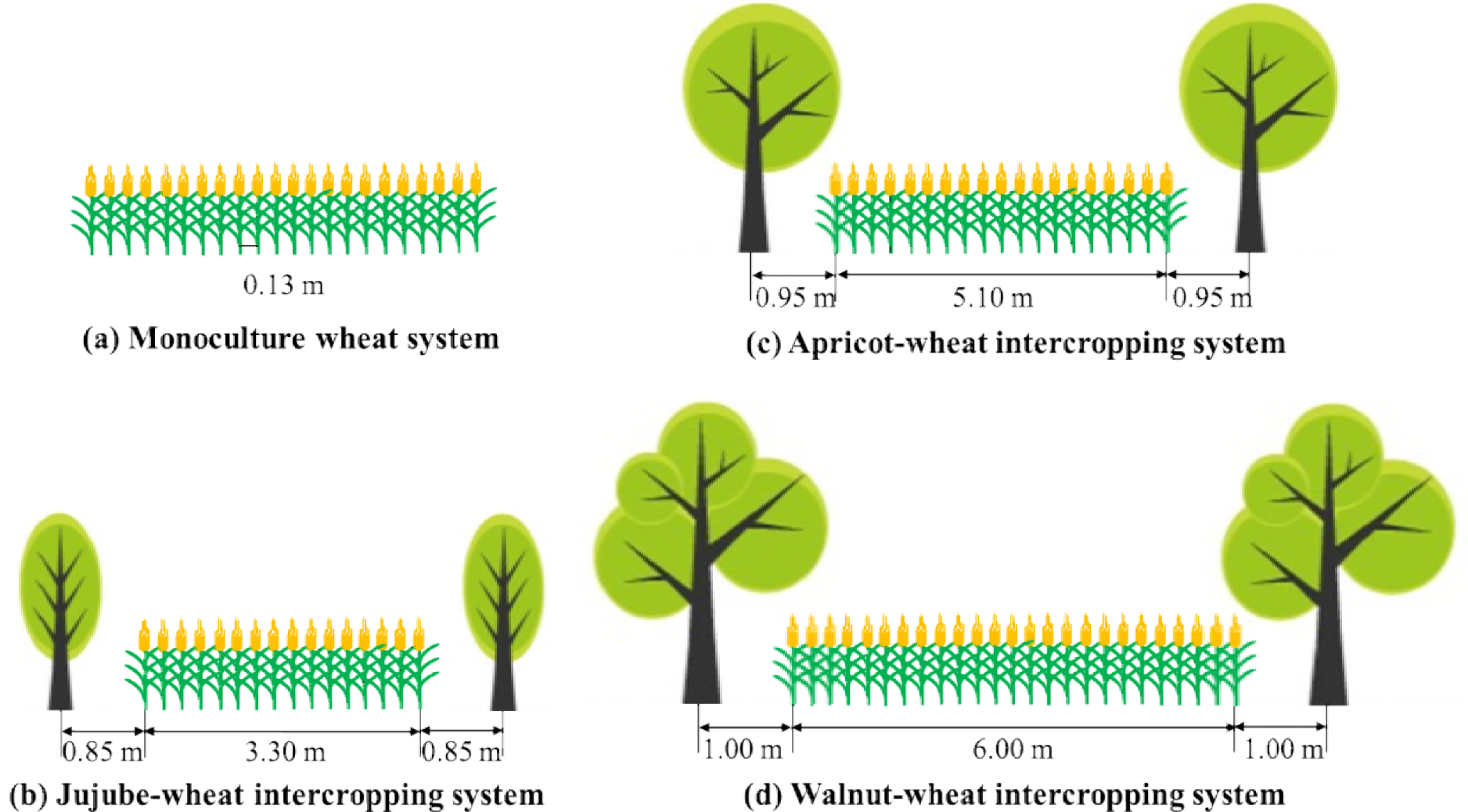
Schematic illustration of planting patterns in monoculture wheat and 3 different fruit tree-wheat based intercropping systems.

In 2011, monoculture wheat and the wheat for the 3 intercropping systems were sown on 8 October, 2010 and harvested on 11 June, 2011. In 2012, the wheat was sown on 3 October, 2011 and harvested on 9 June, 2012. All fields were fertilized with farmyard manure (15,000 kg hm^-2^), urea (275 kg N hm^-2^), triple superphosphate (150 kg P_2_O_5_ hm^-2^), and potassium sulphate (275 kg K hm^-2^); all were applied homogeneously throughout the fields before sowing wheat (40% of the N was applied initially, with the remaining 60% of the N fertilizer applied at the wheat stem elongation stage).

### Harvested and analysis

Wheat was harvested when mature. There were five replicates. In 2011 and 2012, The monoculture wheat, jujube-, apricot-, and walnut-based intercropping wheat were harvested, respectively, 6.5 m^2^(5.0 m length × 1.3 m width), 9.9 m^2^(3.0 m × 3.3 m), 10.2 m^2^(2.0 m×5.1 m), and 15 m^2^(2.5 m × 6.0 m), and samples were immediately dried on a sunning ground to thresh seeds (in order to calculate wheat yield). In order to make wheat samples much more representativeness, 2 m length intercropping wheat samples from three regions (in the middle region of the tree rows, underneath the tree of east canopy and west canopy) were harvested respectively to estimate the total spike number and grains per spike, and then all samples were threshed for seeds to estimate thousand grain weight and harvest index. The stalks (except grains) and grain samples were digested in a mixture of concentrated H_2_SO_4_ and H_2_O_2_. N concentrations were determined using the micro-Kjeldahl method, P concentrations by the molybdo-vanado-phosphate colorimetrical method and K concentrations by flame photometry [18]. In 2011, 15 seedlings for each replication were selected to calculate shoot biomass at overwintering stage, reviving stage, jointing stage, booting stage, anthesis stage, filling stage, and maturity stage, respectively, and the shoot samples were heated at 105 °C for 30 min and then oven-dried (72 h, 75°C) as well.

### Fertile florets

In both years, 21 main spikes from each replicate, which flowering with the same day and the same size, were harvested destructively to investigate fertile florets in the flowering period (50% anthesis). The 21 main spikes were from three regions (in the middle region of the tree rows, underneath the tree of east canopy and west canopy).

### PAR measurement

Light penetration was measured at three regions, in the middle region of the tree rows, under the tree of east canopy and west canopy above wheat using a SunScan Canopy Analysis System (Delta-T Devices, Cambridge, UK). The 64 light sensors of the SunScan measured individual levels of PAR, which are transmitted to a PDA and expressed as μmol·m^-2^·s^-1^. SunScan readings are taken when the sky is clear to avoid the interference of the clouds at the filling stage of wheat in 2011 and 2012. One measurement was performed every two hours from morning at 09:00 until late afternoon at 19:00.

### Photosynthetic parameters

The net photosynthetic rate (Pn) of the flag leaves was determined with a LI-6400XT Portable Photosynthesis System (LI-COR, Inc., USA), and the readings are taken when the sky is clear to avoid the interference of the clouds at the filling stage of wheat in 2011 and 2012. The measurements were conducted under traditional open system and under controlled conditions with a CO_2_ concentration of 380 µmol m^-2^s^-1^. The PAR was set at 1200 µmol m^-2^s^-1^, which was provided by a 6400-2B LED light source. The Pn was measured at three regions, in the middle region of the tree rows, under the tree of east canopy and west canopy. One measurement was performed every two hours from morning at 09:00 until late afternoon at 19:00. An average value was calculated from three flag leaves from each replicate.

### Grain quality analyses

Grain protein content, wet gluten content, falling number, dough development time and dough stability time and softening degree were measured according to official AACC methods [19].

### Statistical analysis

Mean daily shade intensity (%) = (PAR_mono_ - PAR_int_) / PAR_mono_ ×100% Eqn (1) PAR_mono_ is the mean daily PAR of monoculture wheat system; PAR_int_ is the mean daily PAR of fruit tree based intercropping system.

Experimental data was collected from 2011 and 2012. One way analysis of variance was performed on all datasets using SPSS 16.0 for Windows (SPSS Inc., Chicago, IL). Significant differences between pairs of mean values were determined with Duncan’s multiple range test at the 5% level. Standard error between the replications was also calculated. Simple regression analysis was used to examine the relationships among the data of fertile florets, grain yield (including spike number, grains per spike, and thousand grain weight), nutrient uptake (N, P, and K content) and grain quality (including protein content, wet gluten content, falling number, dough development time, dough stability time, and softening degree) of wheat with understory mean daily shade intensity.

## Results

### Light interception and photosynthetic rate

Diurnal variation of the understory photosynthetically active radiation (PAR) and photosynthetic rate (Pn) in the three intercropping systems and the monoculture wheat system varied with time, and with single peak curves during midday (13:00–15:00) (Fig 2). Owing to reflectance, absorbance, and transmittance by the canopies of the three fruit tree types, the PAR of crops in the intercropping systems were lower than that in the monoculture configurations. For example, the mean daily PAR in the jujube-, apricot-, and walnut-based intercropping systems were, respectively, just 78.7%, 45.5%, and 20.1% of the monoculture configurations in 2011 and 75.0%, 39.5%, and 18.9% of the monoculture configurations in 2012 (Fig 2a, b). Further, the photosynthetic rates in the jujube-, apricot-, and walnut-based intercropping systems decreased, respectively, by an average 26.2%, 36.9%, and 50.9% compared to monoculture wheat in 2011 and by 26.6%, 37.9%, and 48.2% in 2012.

**Figure 2.**
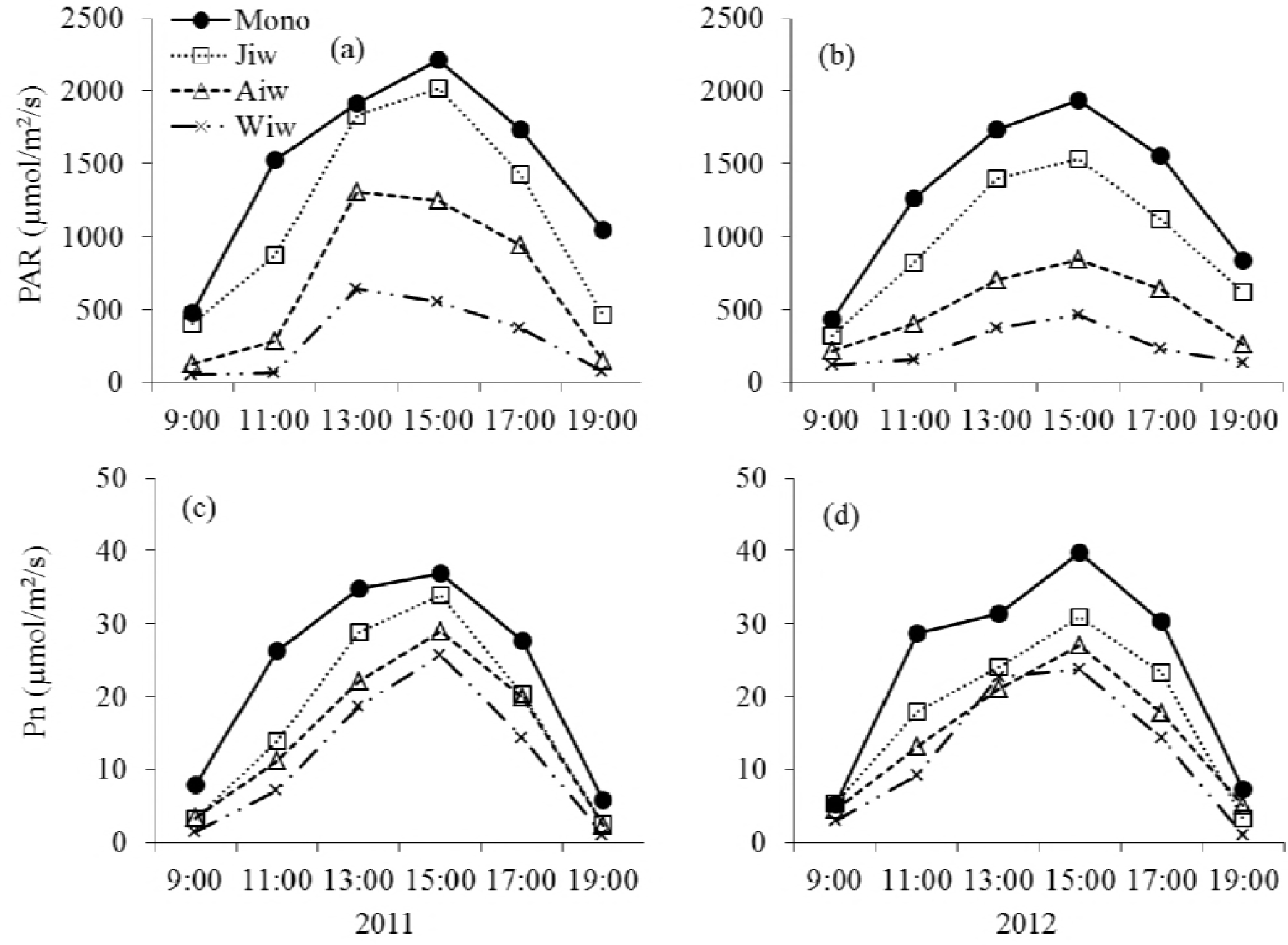
The daily change of photosynthetically active radiation (PAR) and photosynthetic rate (Pn) of wheat in monoculture configurations and 3 different fruit tree-wheat intercropping systems at the filling stage in 2011 (a, c) and 2012 (b, d). The PAR and Pn data are the mean values of the three regions, in the middle region of the tree rows, under the tree of east canopy and west canopy, respectively. Mono, monoculture wheat system; Jiw, jujube-wheat intercropping system; Aiw, apricot-wheat intercropping system; Wiw, walnut-wheat intercropping system.

### Fertile florets

The distribution of fertile florets along the wheat spikes is shown in Figure 3. Fruit tree shade reduced the number of fertile florets on almost all spikelets, with especially pronounced reductions in the middle position (spikelets 4-12 from the base of the spike). The total number of fertile florets per wheat spike in the monoculture configuration were increased by 1.12, 1.35, and 1.42 times compared to the jujube-, apricot-, and walnut-based intercropping systems in 2011 and by 1.14, 1.61, and 1.76 times in 2012, respectively (Fig 3). Furthermore, significant correlations (*P* < 0.001) were observed between the number of fertile florets and the mean daily shade intensity of wheat in both 2011 and 2012 (Fig 4).

**Figure 3.**
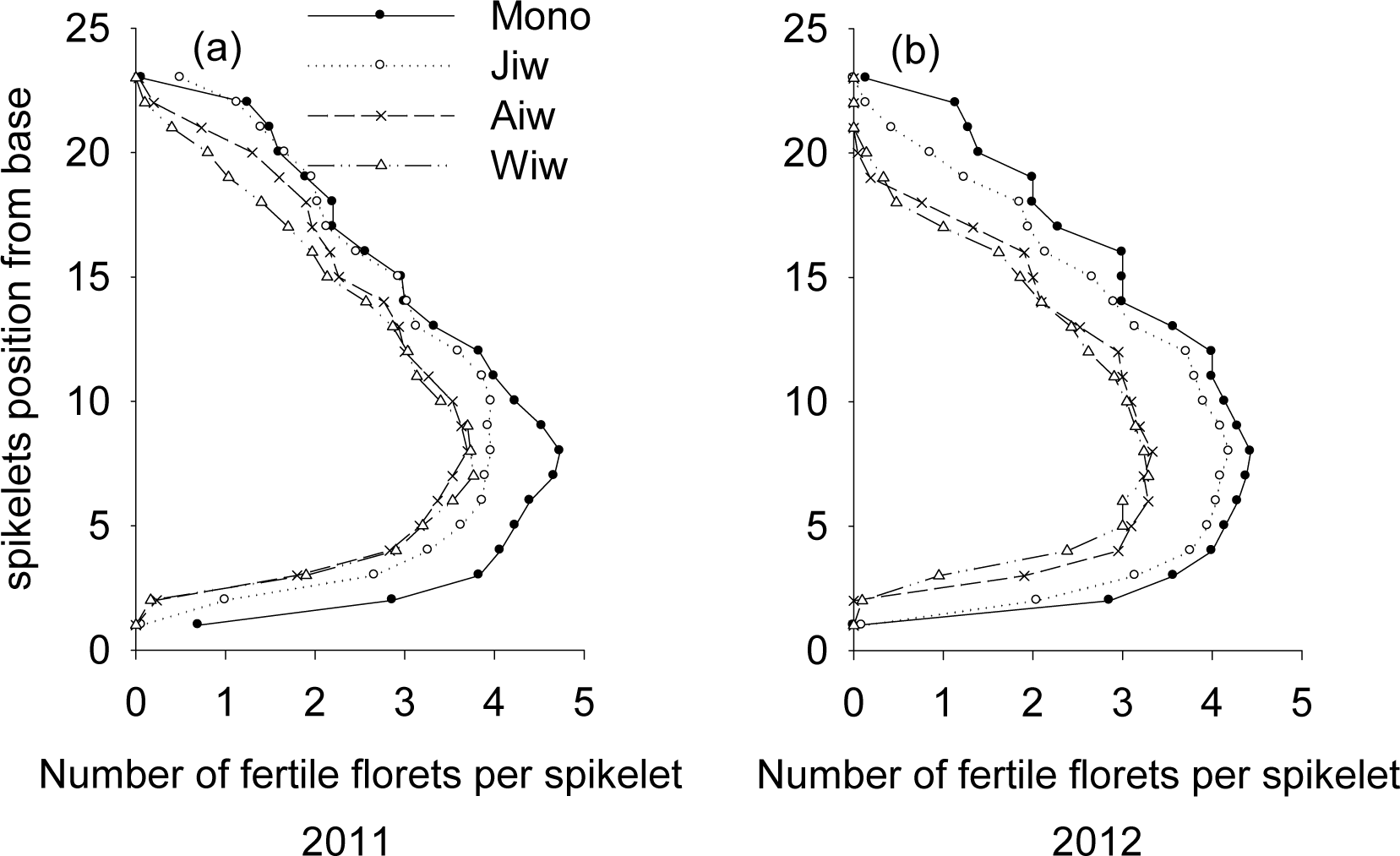
Distribution of the fertile florets along the spike of wheat in monoculture configurations and 3 different fruit tree-wheat intercropping systems in 2011 (a) and 2012 (b). The distribution of the fertile florets data are the mean values of the three regions, in the middle region of the tree rows, under the tree of east canopy and west canopy. Mono, monoculture wheat system; Jiw, jujube-wheat intercropping system; Aiw, apricot-wheat intercropping system; Wiw, walnut-wheat intercropping system.

**Figure 4.**
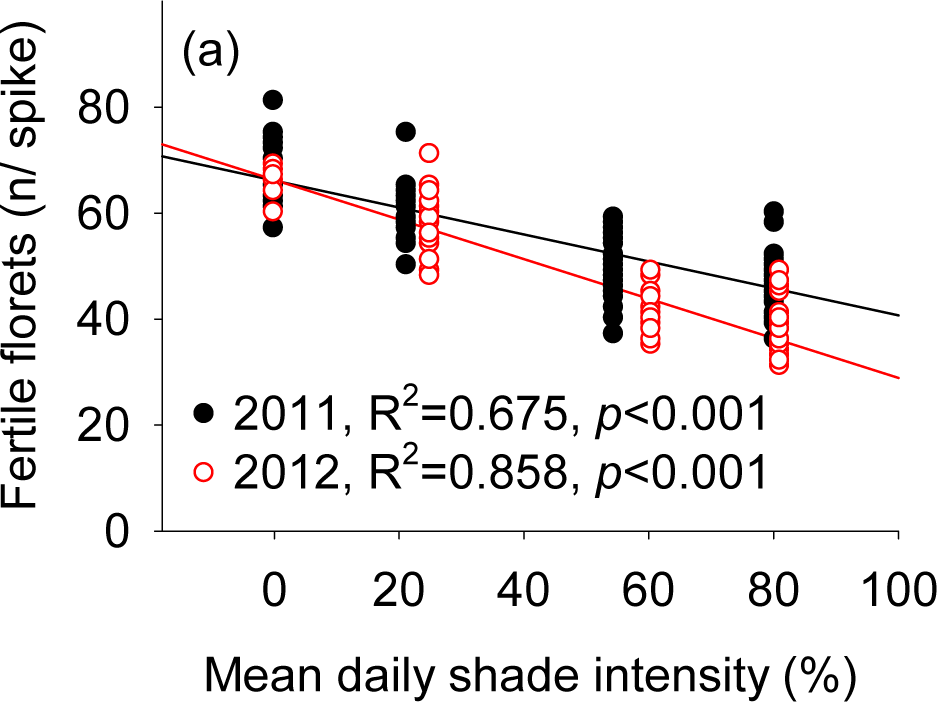
Relationship between the fertile florets and mean daily shade intensity in 2011 and 2012.

### Wheat yield components

In 2011 and 2012, spike number (expressed per unit area of the monoculture wheat or the real intercropping wheat strip area—*i.e.*, without the distance from the fruit trees to the nearest wheat row) and grains per spike were significantly higher in the monoculture wheat and jujube-based intercropping wheat systems than in the apricot- and walnut-based intercropping systems (Table 1). In both years, the thousand grain weight, harvest index (proportion of seed dry weight relative to the total above-ground dry weight), and net yield of wheat in the monoculture wheat system were each significantly higher than in the jujube-, apricot-, and walnut-based intercropping systems (excluding the net yield of wheat in the jujube-based intercropping system in 2011). Additionally, in both 2011 and 2012, strong negative linear correlations (*P* < 0.001) were observed between mean daily shade intensity and spike number, grains per spike, thousand grain weight, and net yield (Fig 5).

**Table 1.**
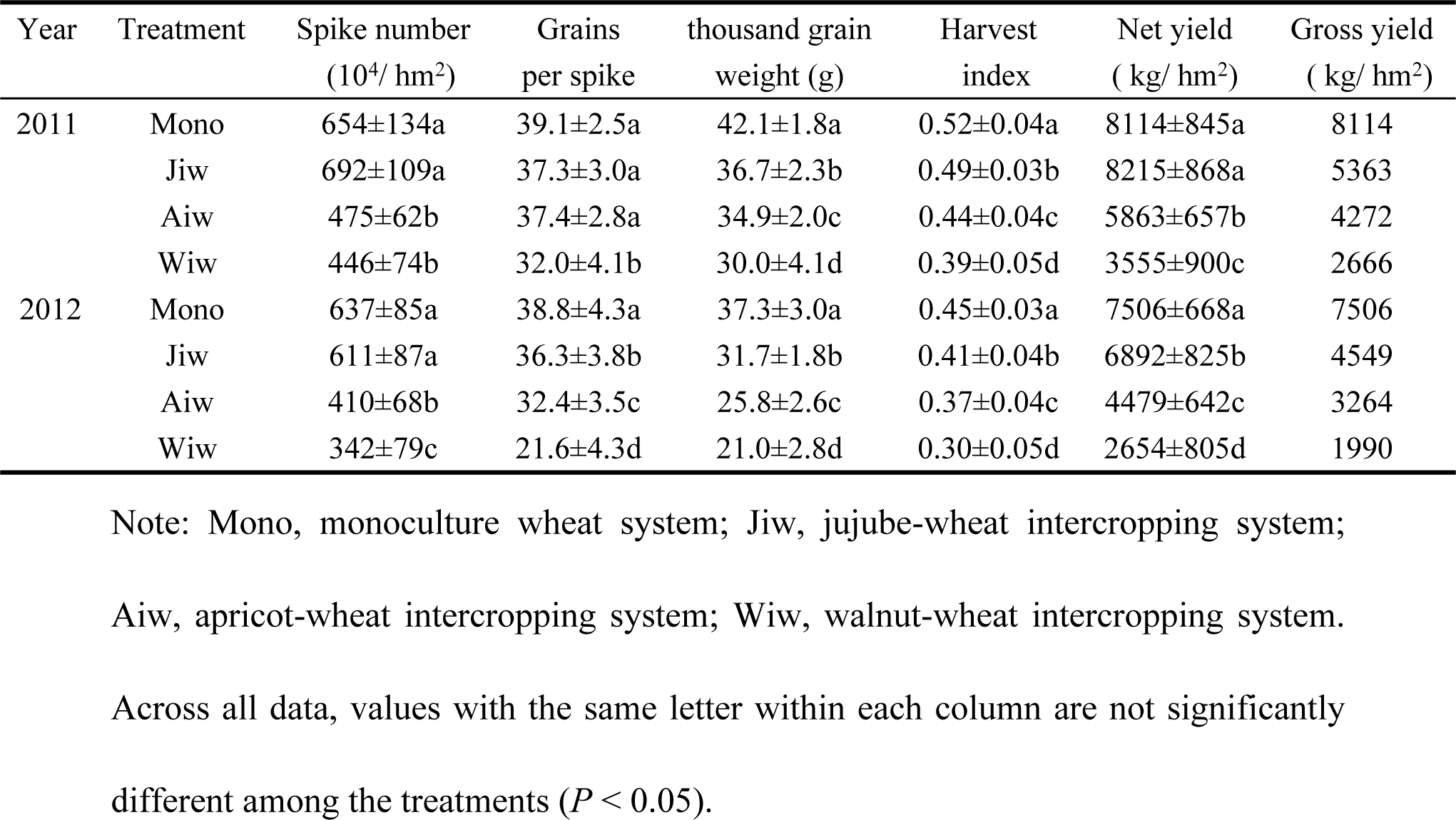
Yield components of wheat in monoculture configurations and 3 different fruit tree-wheat intercropping systems in 2011 and 2012.

| Year | Treatment | Spike number<br>(10<sup>4</sup>/ hm<sup>2</sup>) | Grains<br>per spike | thousand grain<br>weight (g) | Harvest<br>index | Net yield<br>( kg/ hm<sup>2</sup>) | Gross yield<br>( kg/ hm<sup>2</sup>) |
| --- | --- | --- | --- | --- | --- | --- | --- |
| 2011 | Mono | 654±134a | 39.1±2.5a | 42.1±1.8a | 0.52±0.04a | 8114±845a | 8114 |
|  | Jiw | 692±109a | 37.3±3.0a | 36.7±2.3b | 0.49±0.03b | 8215±868a | 5363 |
|  | Aiw | 475±62b | 37.4±2.8a | 34.9±2.0c | 0.44±0.04c | 5863±657b | 4272 |
|  | Wiw | 446±74b | 32.0±4.1b | 30.0±4.1d | 0.39±0.05d | 3555±900c | 2666 |
| 2012 | Mono | 637±85a | 38.8±4.3a | 37.3±3.0a | 0.45±0.03a | 7506±668a | 7506 |
|  | Jiw | 611±87a | 36.3±3.8b | 31.7±1.8b | 0.41±0.04b | 6892±825b | 4549 |
|  | Aiw | 410±68b | 32.4±3.5c | 25.8±2.6c | 0.37±0.04c | 4479±642c | 3264 |
|  | Wiw | 342±79c | 21.6±4.3d | 21.0±2.8d | 0.30±0.05d | 2654±805d | 1990 |

Note: Mono, monoculture wheat system; Jiw, jujube-wheat intercropping system;
Aiwi, apricot-wheat intercropping system; Wiw, walnut-wheat intercropping system.
Across all data, values with the same letter within each column are not significantly
different among the treatments ( $P < 0.05$ ).

**Figure 5.**
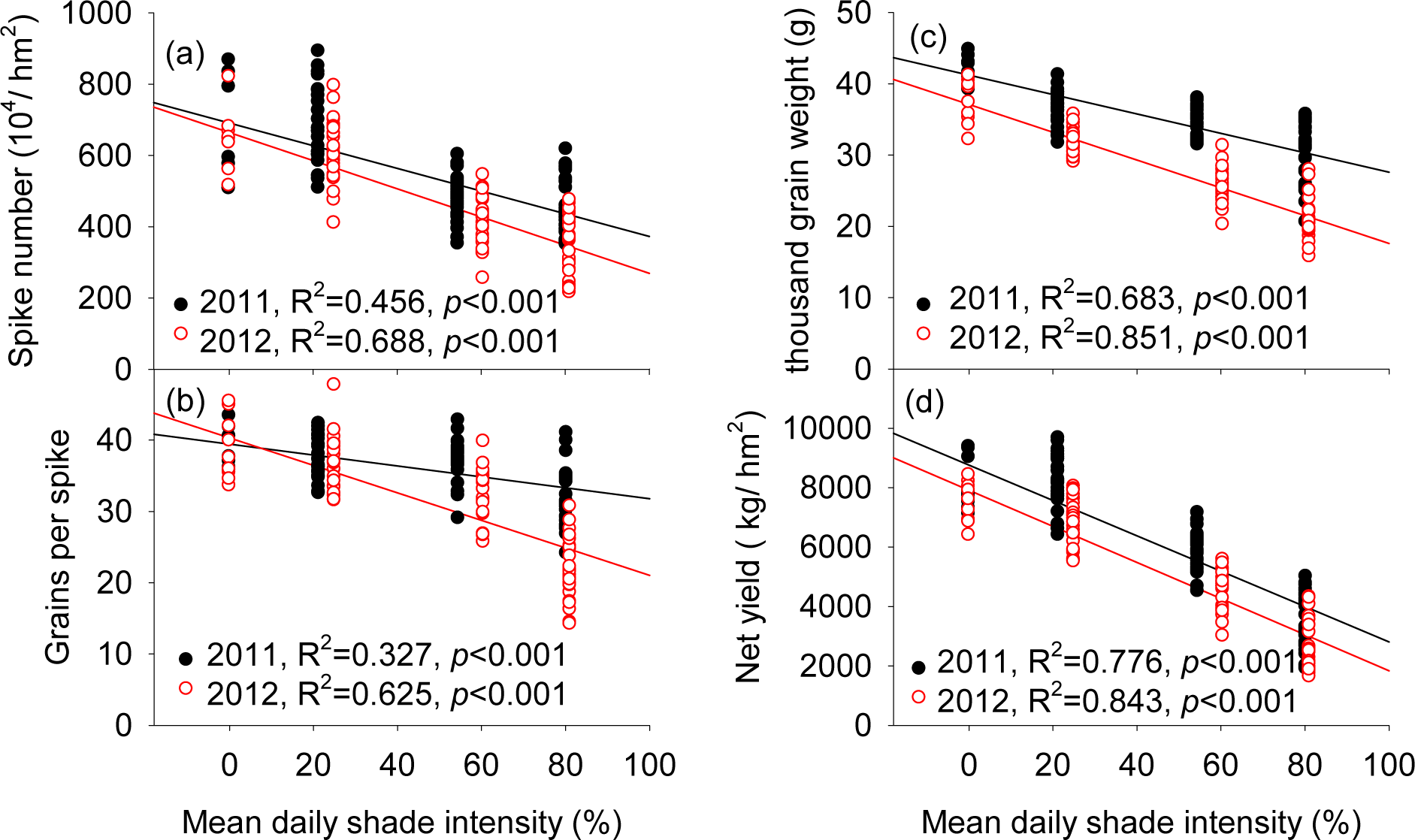
Relationship between the spike number (a), grains per spike (b), thousand grain weight (c) and net yield (d) with mean daily shade intensity of wheat in 2011 and 2012.

### N, P, and K content

In 2011 and 2012, the total N, P, and K uptake of wheat in the monoculture system and the jujube-based intercropping systems were significantly higher than in the walnut-based intercropping system (Table 2). For example, the N, P, and K content of wheat in the monoculture system were, respectively, 1.55, 1.63, and 1.50 times higher than in the walnut-based intercropping system in 2011 and 1.56, 1.75 and 1.61 times higher in 2012. Additionally, in both 2011 and 2012, strong negative linear correlations (*P* < 0.01) were observed between mean daily shade intensity of wheat and N, P, and K content (Fig 6).

**Table 2.** The N, P and K contents of wheat in monoculture configurations and 3 different fruit tree-wheat intercropping systems in 2011 and 2012.

259 different fruit tree-wheat intercropping systems in 2011 and 2012.
| Year | Treatment | N content (kg/ hm<sup>2</sup>) | P content (kg/ hm<sup>2</sup>) | K content (kg/ hm<sup>2</sup>) |
| --- | --- | --- | --- | --- |
| 2011 | Mono | 191±9a | 29.9±3.4a | 548±34a |
|  | Jiw | 198±8a | 29.4±2.0a | 583±47a |
|  | Aiw | 169±8b | 24.3±2.7a | 519.6±23a |
|  | Wiw | 138±19c | 18.3±3.4b | 366±60b |
| 2012 | Mono | 189±7a | 29.4±0.3a | 607±13a |
|  | Jiw | 195±11a | 28.2±3.3a | 604±73a |
|  | Aiw | 169±16b | 20.7±1.8b | 473±16b |
|  | Wiw | 121±5c | 16.8±1.2c | 377±35c |

Note: Mono, monoculture wheat system; Jiw, jujube-wheat intercropping system; Aiw, apricot-wheat intercropping system; Wiw, walnut-wheat intercropping system. Across all data, values with the same letter within each column are not significantly different among the treatments ( $P < 0.05$ ).

**Figure 6.**
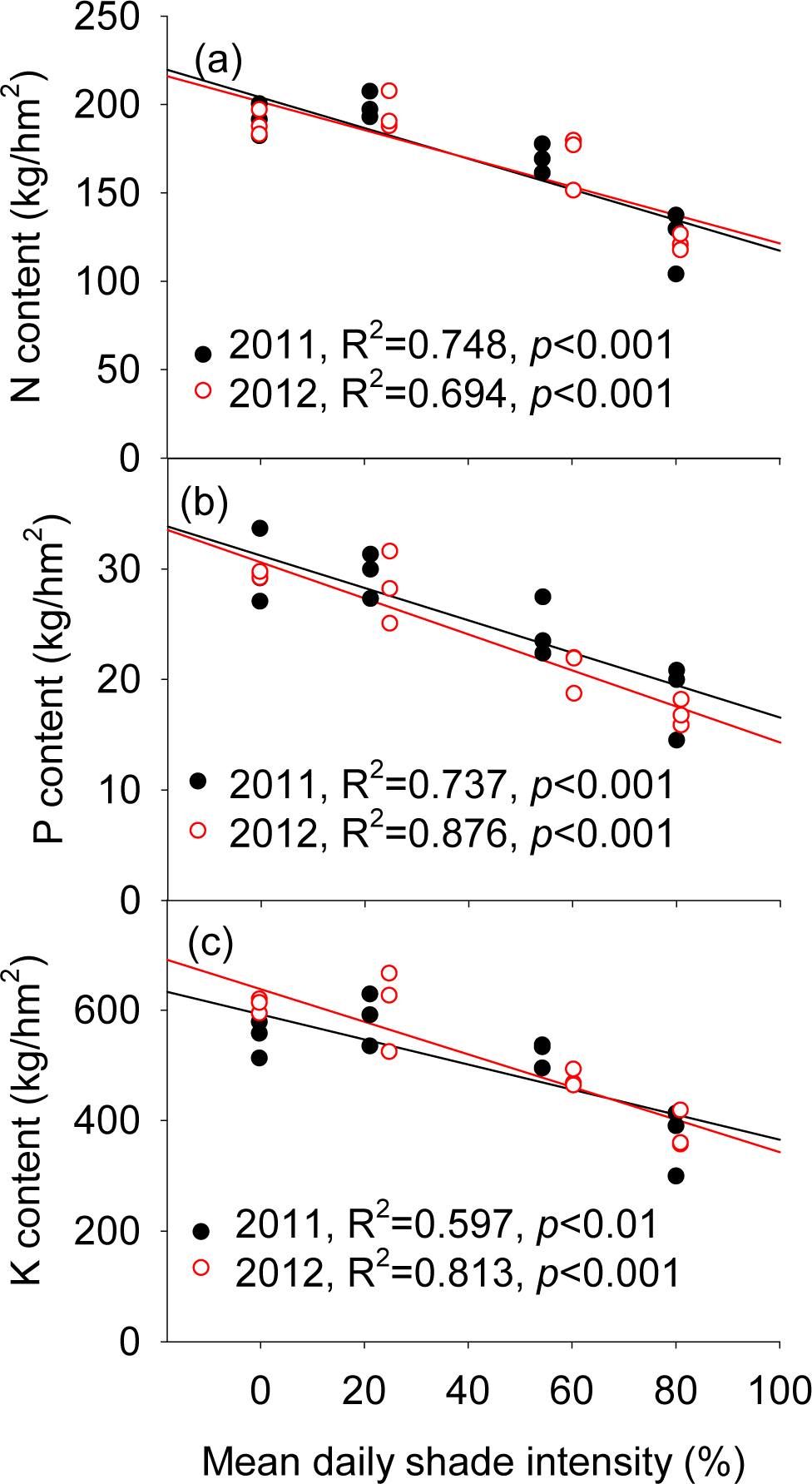
Relationship between the N, P and K contents with mean daily shade intensity of wheat in 2011 and 2012.

### Grain quality traits

In 2011 and 2012, protein content, wet gluten content, falling number, dough development time, and dough stability time of wheat in the walnut-based intercropping system were significantly higher than in the monoculture and jujube-based intercropping system. In contrast, the highest values for the softening degree parameter were observed in the monoculture system (Table 3). Furthermore, the mean daily shade intensity of wheat both in 2011 and 2012 was highly positively linearly correlated (*P* < 0.01) with protein content, wet gluten content, falling number, development time, and stability time (Fig 7a, b, c, d & e). The softening degree was negatively linearly correlated (*P* < 0.01) with mean daily shade intensity (Fig 7f).

**Table 3.**
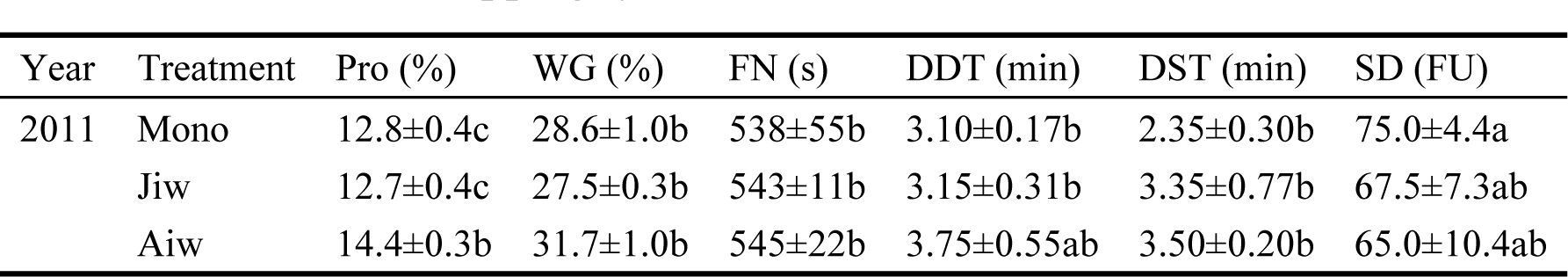
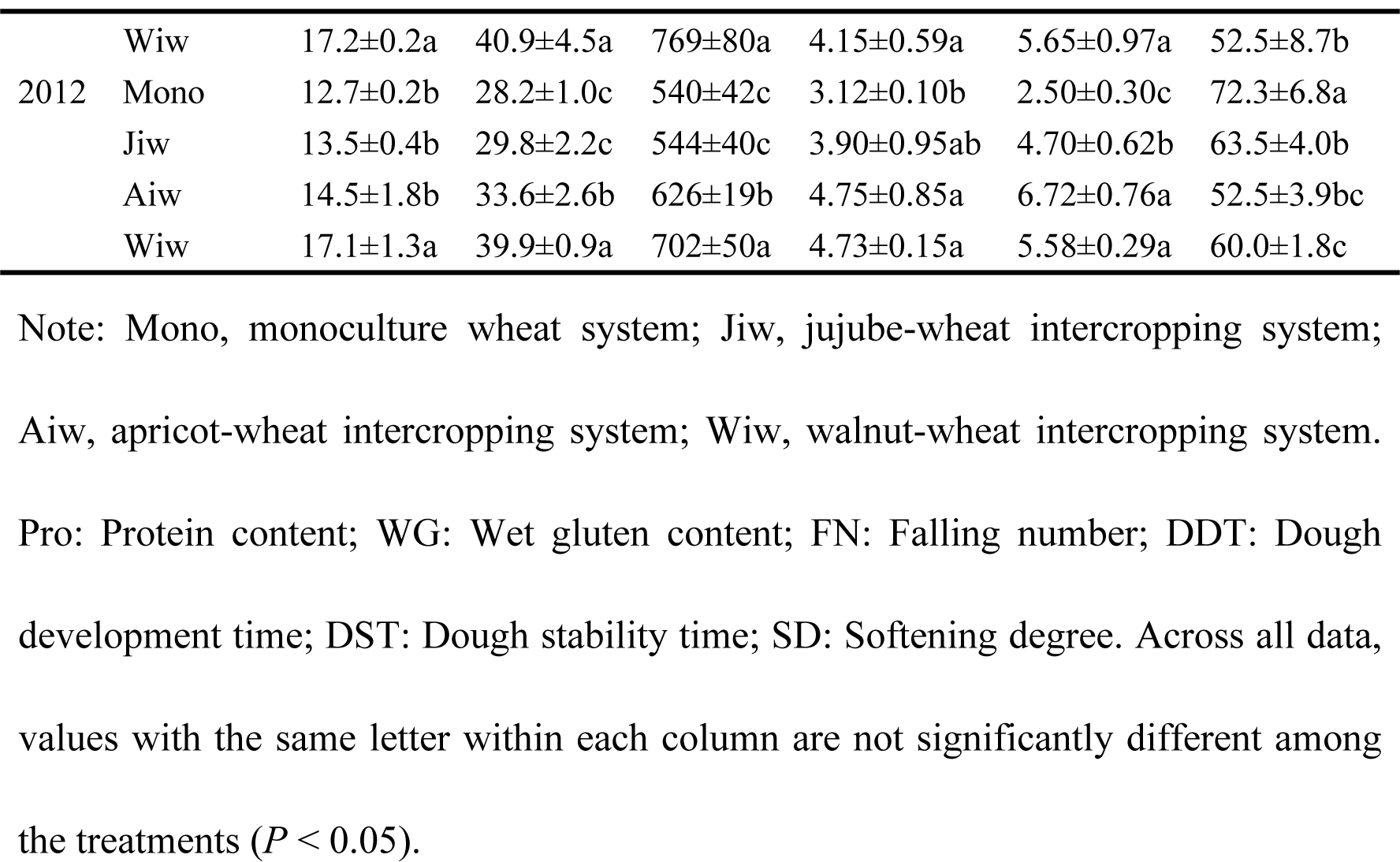
the grain quality of wheat in monoculture configurations and 3 different fruit tree-wheat intercropping systems in 2011 and 2012.

| Year | Treatment | Pro (%) | WG (%) | FN (s) | DDT (min) | DST (min) | SD (FU) |
| --- | --- | --- | --- | --- | --- | --- | --- |
| 2011 | Mono | 12.8±0.4c | 28.6±1.0b | 538±55b | 3.10±0.17b | 2.35±0.30b | 75.0±4.4a |
|  | Jiw | 12.7±0.4c | 27.5±0.3b | 543±11b | 3.15±0.31b | 3.35±0.77b | 67.5±7.3ab |
|  | Aiw | 14.4±0.3b | 31.7±1.0b | 545±22b | 3.75±0.55ab | 3.50±0.20b | 65.0±10.4ab |
|  | Wiw | 17.2±0.2a | 40.9±4.5a | 769±80a | 4.15±0.59a | 5.65±0.97a | 52.5±8.7b |
| 2012 | Mono | 12.7±0.2b | 28.2±1.0c | 540±42c | 3.12±0.10b | 2.50±0.30c | 72.3±6.8a |
|  | Jiw | 13.5±0.4b | 29.8±2.2c | 544±40c | 3.90±0.95ab | 4.70±0.62b | 63.5±4.0b |
|  | Aiw | 14.5±1.8b | 33.6±2.6b | 626±19b | 4.75±0.85a | 6.72±0.76a | 52.5±3.9bc |
|  | Wiw | 17.1±1.3a | 39.9±0.9a | 702±50a | 4.73±0.15a | 5.58±0.29a | 60.0±1.8c |

Note: Mono, monoculture wheat system; Jiw, jujube-wheat intercropping system; Aiwi, apricot-wheat intercropping system; Wiw, walnut-wheat intercropping system. Pro: Protein content; WG: Wet gluten content; FN: Falling number; DDT: Dough development time; DST: Dough stability time; SD: Softening degree. Across all data, values with the same letter within each column are not significantly different among the treatments ( $P < 0.05$ ).

**Figure 7.**
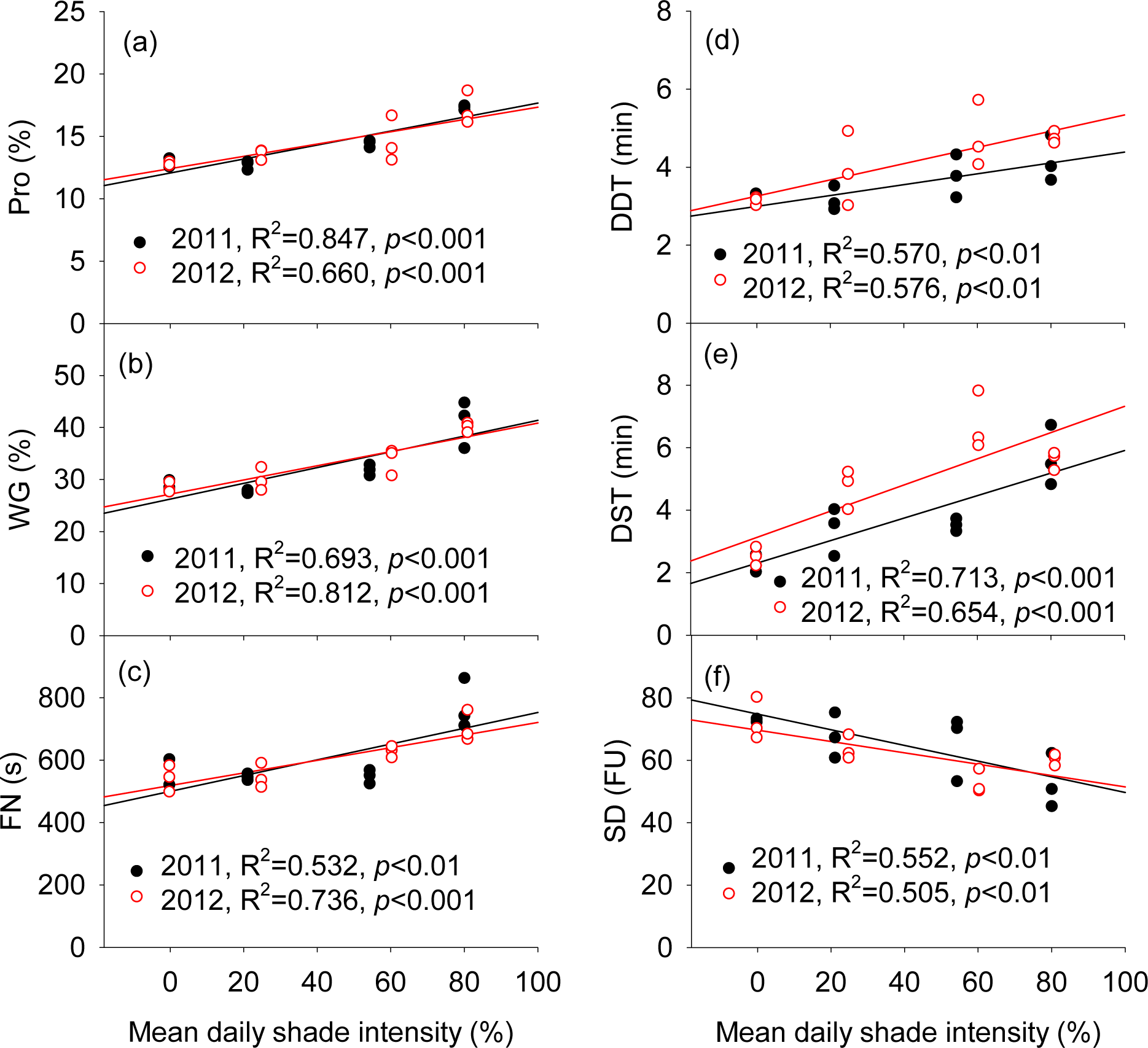
Relationships between grain quality with mean daily shade intensity of wheat in 2011 and 2012. Pro: Protein content; WG: Wet gluten content; FN: Falling number; DDT: Dough development time; DST: Dough stability time; SD: Softening degree.

## Discussion

### PAR and photosynthetic rate

Light, as a primary limiting factor in tree-based intercropping systems, influences the growth and development of intercropped crops significantly [20]. Awal et al. [21] and Reynolds et al. [22] showed that it was difficult for maize to obtain sufficient solar energy when grown underneath the canopy of higher jujube tree components at a distance of less than 2.5 m to the tree rows. Similar results were reported in other studies of temperate agroforestry systems [12,23,24]. In our study, the mean daily shade intensity in jujube-, apricot-, and walnut-based intercropping systems compared to monoculture configurations was 21.3%, 54.5%, and 80.3% shade, respectively, in 2011, and 25.0%, 60.5%, and 81.1% in 2012 (Fig 2). Gao et al. [15] observed a clear, positive linear relationship between the distance from the apple tree rows and the daily mean values of PAR and net photosynthetic rate in apple–soybean and apple– peanut intercropping systems. Additionally, Kittur et al. [12] reported that the rhizome yield and understory PAR could be predicted by a linear equation in Kerala, India. In our study, in the walnut tree-based intercropping system, due to its taller trunk, larger leaves, and large canopy architecture, the intercropped wheat was markedly influenced by the shading effect of the trees. The photosynthetic rate of walnut-based intercropped wheat was 50.9% and 48.2% lower than that of monoculture wheat in 2011 and 2012, respectively. Regular pruning of the fruit-tree canopy could reduce light interception, thus improving the yield of intercropped crops.

### Fertile florets

The grain number per spike is determined by the number of spikelets per spike and the number of florets per spikelet. The environmental factors (e.g., light) that determine the spikelet number of grains have been well studied [25]. In the present study, the fruit tree-based intercropping system reduced the number of fertile wheat florets on almost all spikelets, particularly in the middle portions of 4–12 spikelets of the spike (Fig 3). Therefore, the development of the floret varies considerably depending on its position on the spike [25]. Furthermore, we observed significant (*P* < 0.001), negative correlations between the fertile florets and mean daily shade intensity from the fruit trees (Fig 4). Willey and Holliday [26] showed that light intensity significantly influenced the initiation of spikelets and floret primordia, giving rise to fewer grains. For example, shading delayed the rate of floret initiation per spike by 11.4%, consequently decreasing the number of florets by 22.3% and the grain weight per spike by 19% at maturity [25].

### Grain yield components

Understory crop yield is determined by the intercepted available light, and the efficiency of converting the intercepted light into photosynthate [15]. Peng et al. [14] reported decreases of 38% and 29% in the yields of maize and soybean, respectively, in a tree-based agroforestry intercropping system on the Loess Plateau, China. In our study, spike number, grains per spike, thousand-grain weight, and net yield were all significantly higher in the monoculture wheat and jujube-based intercropping wheat systems than those in the apricot- and walnut-based intercropping systems in both 2011 and 2012 (Table 1). Additionally, the wheat shoot biomass in the apricot- and walnut-based intercropping systems was 21.4% and 42.5% lower, respectively, at the mature stage in 2011 (S1 Fig.). Yang et al. [27] reported that the yield of intercropped wheat was decreased by 25.8%, 16.5%, and 6.70% at distances of 90, 110, and 130 cm to the jujube tree rows, respectively. Other studies of temperate agroforestry systems reported significant increases in the grain yield and yield components with increasing distance to the tree rows [14,28]. We observed a highly significant (*P* < 0.001), negative linear correlation between the wheat grain yield and its components (spike number, grains per spike, and thousand-grain weight) and mean daily shade intensity (Fig 5). Thus, our study demonstrates that the shading intensity of fruit trees in the agroforestry systems had a significant, negative effect on the intercropped grain yield and its components.

### Shoot nitrogen (N), phosphorous (P), and potassium (K) uptake

Light plays an important role in dry matter accumulation and the nutrient uptake of crops. Cui et al. [29] showed that shade significantly decreased the total N, P, and K contents of summer maize. In the present study, the total N, P, and K contents of wheat in the walnut-based intercropping systems were significantly lower than those in monoculture wheat (Table 2). Previous studies showed that nutrient uptake by the understory crop were closely correlated with overstory tree density, the understory plant varieties, and plant nutrient demand [15,30]. Kittur et al. [12] reported decreased uptake of N, P, and K by understory turmeric with decreasing bamboo spacing. We observed a highly significant (*P* < 0.05), negative linear correlation between the N, P, and K contents of wheat and mean daily shade intensity (Fig 6). The N, P, and K concentrations of wheat stalks and grains in the walnut-based intercropping systems were significantly higher than those in monoculture wheat (S3 Table). Cui et al. [29] also reported that the N, P, and K concentrations of summer maize were enhanced by shading. The fruit-tree shade intensity increased the dry matter accumulation of the intercropped wheat, which in turn increased the ability of the wheat plants to absorb nutrients.

### Grain quality

Lu et al. [31] reported that the protein and wet gluten contents, and the falling number of intercropped wheat increased significantly in a *Paulownia*-based intercropping system compared to a monoculture configuration. Additionally, Wang et al. [32] reported that the wheat starch and crude fat contents were enhanced in apricot tree-based agroforestry systems, and that the quality of the wheat decreased with decreasing distance to the apricot trees. In the present study, we found that increased tree shade intensity markedly enhanced the wheat protein and wet gluten contents, falling number, and dough development and stability times, whereas it significantly decreased the degree of softening (Table 3). Our results are generally consistent with those from previous studies and confirm that tree shading during grain development can have a substantial effect on grain yield and quality in agroforestry systems [32]. We observed a highly significant (*P* < 0.01), positive linear correlation between the wheat protein and wet gluten contents, falling number, and dough development and stability times with the mean daily shade intensity, whereas shade was negatively correlated with the degree of softening (Fig 7). Bhatta et al. [33] reported a negative correlation between grain protein content and grain yield and grain volume weight. In our study, we observed a significant, positive linear correlation between the protein and wet gluten contents, falling number, and dough development and stability times and the grain yield and thousand-grain weight of wheat, and a negative correlation with the degree of softening (S2 and S3 Figs.). Consequently, fruit tree shading resulted in a decrease in wheat yields and seed dry weight, resulting in a decrease in the quality of wheat.

Friday and Fownes [11] suggested that the shade intensity from trees could be alleviated by pruning of the tree canopy, increasing the intercepted light reaching the intercropped plants. To obtain higher grain production, appropriate management measures are needed to minimize competition in fruit tree–crop intercropping systems, and we recommend the following measures: (1) select more suitable crop varieties (e.g., shade-tolerant, early-maturing, and low-height varieties); (2) select more suitable fruit tree varieties (e.g., low-height varieties that have small leaves); and (3) regularly prune the fruit trees to reduce the shading intensity from the fruit trees.

## Conclusion

Tree-based intercropping systems are a common traditional practice in China, owing to their significant favorable effects on reducing soil and water losses, increasing land-use efficiency, increasing economic returns, and improving ecological environments, and these systems are particularly important in southern Xinjiang. We found that tree shade intensity was generally the major limiting factor for crop productivity in agroforestry systems in this region. Reflectance, absorbance, and transmittance by the tree canopy dramatically reduce the PAR for the crop canopy. Fruit-tree shading resulted in decreased photosynthate accumulation, which in turn resulted in decreased nutrient uptake in the intercropped grain plants. Fruit-tree shading had a marked, negative effect on the development of fertile florets, resulting in fewer grains per spike and reduced net photosynthesis, ultimately resulting in decreases in grain weight, grain yield, and grain quality. Our results show that jujube-based intercropping systems offer a suitable agroforestry system in the region since they did not decrease the yield and quality of the intercropped wheat. Highly significant, negative linear correlations were observed between tree shade intensity and the number of fertile florets, grain yield (including spike number, grains per spike, and thousand-grain weight), nutrient content (N, P, and K), and softening degree of wheat. In contrast, we found that daily shade intensity was positively linearly correlated with wheat protein content, wet gluten content, falling number, and the dough development and stability times. Future research should focus on the development of shade-tolerant crop varieties and examine how regular pruning of the tree canopy structure can improve crop productivity in such systems. It would also be useful to investigate the mechanisms underlying the aboveground and belowground interspecific interactions in agroforestry systems further.

## Supporting Information

**S1 Table. Nutrient status of the experimental soil.**

**S2 Table. The basic information of the 3 fruit trees**

**S3 Table. N, P and K concentrations in stalks and grains of wheat in monoculture configurations and 3 different fruit tree-wheat intercropping systems in 2011 and 2012.**

**S1 Fig. The shoot dry weight of wheat in monoculture configurations and 3 different fruit tree-wheat intercropping systems in 2011.**

**S2 Fig. Relationships between grain quality and grain net yield of wheat in 2011 and 2012.**

**S3 Fig. Relationships between grain quality and thousand grain weight of wheat in 2011 and 2012.**

## Acknowledgments

We are grateful for the support from the Extension Centre of Agricultural Technology in Zepu County, Kashi Prefecture, Xinjiang. We also would like to thank the anonymous reviewers and the editors for their helpful comments.

## Author Contributions

XQ and XC designed the research; XQ, LS and LX performed the research; XQ and JL analyzed the data; and XQ and JL wrote the paper.

